# AAV-mediated Delivery of Plakophilin-2a Arrests Progression of Arrhythmogenic Right Ventricular Cardiomyopathy in Murine Hearts: Preclinical Evidence Supporting Gene Therapy in Humans

**DOI:** 10.1101/2023.07.12.548590

**Authors:** Chantal JM van Opbergen, Bitha Narayanan, Chester B Sacramento, Katie M Stiles, Vartika Mishra, Esther Frenk, David Ricks, Grace Chen, Mingliang Zhang, Paul Yarabe, Jonathan Schwartz, Mario Delmar, Chris D Herzog, Marina Cerrone

**Author notes:** These authors contributed equally to the work (Chantal JM van Opbergen, Bitha Narayanan). Equally contributed (Mario Delmar, Chris D Herzog, Marina Cerrone). Co-corresponding authors. Correspondence: Marina Cerrone, MD, Research Associate Professor of Medicine Leon H. Charney Division of Cardiology, New York University Grossmann School of Medicine Science Building 435 E.30^th^ Street 723H, Mario Delmar, MD, PhD, Patricia M. and Robert H. Martinsen Professor of Medicine Leon H. Charney Division of Cardiology, New York University Grossmann School of Medicine Science Building 435 E.30^th^ Street 707, Christopher D. Herzog, Ph.D. Assoc Vice President, AAV R&D Rocket Pharmaceuticals, Inc., 9 Cedar Brook Drive Cranbury, NJ 08512.

## Abstract

**Background:** Pathogenic variants in plakophilin-2 (PKP2) cause arrhythmogenic right ventricular cardiomyopathy (ARVC), a disease characterized by life-threatening arrhythmias and progressive cardiomyopathy leading to heart failure. No effective medical therapy is available to prevent and/or arrest the disease. We tested the hypothesis that AAV-mediated delivery of the human *PKP2* gene to an adult mammalian heart deficient in PKP2 can arrest disease progression and significantly prolong survival.

**Methods:** Experiments were carried out using a cardiac-specific, tamoxifen (TAM)-activated PKP2 knockout murine model (PKP2-cKO). The potential therapeutic, AAVrh.74-PKP2a (RP-A601), is a recombinant AAVrh.74 gene therapy viral vector encoding the human PKP2 variant A (PKP2a). AAVrh.74-PKP2a was delivered to adult mice by a single tail vein injection either before or after TAM-activated PKP2-cKO. PKP2 expression was confirmed by molecular and histopathologic analyses. Cardiac function and disease progression were monitored by survival analyses, echocardiography and electrocardiography.

**Results:** Consistent with prior findings, loss of PKP2 expression caused 100% mortality within 50 days after TAM injection. In contrast, AAVrh.74-PKP2a-mediated PKP2a expression resulted in 100% survival for more than 5 months (at study termination). Echocardiographic analysis revealed that AAVrh.74-PKP2a prevented right ventricle dilation, arrested left ventricle functional decline, and mitigated arrhythmia burden. Molecular and histological analysis showed AAVrh.74-PKP2a– mediated transgene mRNA and protein expression and appropriate PKP2 localization at the cardiomyocyte intercalated disc. Importantly, therapeutic benefit was shown in mice receiving AAVrh.74-PKP2a *after* disease onset.

**Conclusion:** These preclinical data demonstrate the potential for AAVrh.74-PKP2a (RP-A601) as a therapeutic for PKP2-related ARVC in both early and more advanced stages of disease.

## INTRODUCTION

Pathogenic variants in the *PKP2* gene, coding for the protein plakophilin-2 (PKP2) are the primary cause of gene-positive arrhythmogenic right ventricular cardiomyopathy (ARVC) in humans, a progressive disorder with autosomal dominant inheritance.^1, 2^ ARVC is a condition that falls within the umbrella term of arrhythmogenic cardiomyopathy (ACM), as defined in Towbin et al.^3^ For simplicity, in the present document we refer to PKP2-related ARVC as “PKP2-ACM.” Average age at first presentation is estimated between the second to fourth decade and the disease is symptomatic more commonly in males.^2, 4^ Pathogenic variants in PKP2 account for 20-45% of all cases of arrhythmogenic cardiomyopathy cases, with an estimated prevalence between 1:1000 to 1:5000 in Europe and North America.^1^ ARVC is characterized by a high risk of life-threatening arrhythmias prior to other clinical manifestations (i.e., during the concealed phase of the disease), followed by a stage of clinically overt loss of myocardial mass and the presence of fibrofatty infiltrates.^2^ The cardiomyopathy phenotype presents first in the right ventricle, later progressing to a biventricular disease.^1, 2^ ARVC is a disease without a cure and, when advancing to end-stage failure, death can only be prevented by a cardiac transplant.

AAVrh.74-PKP2a (RP-A601) is a recombinant AAVrh.74 gene therapy viral vector encoding the human plakophilin 2 transcript variant A (PKP2a) being developed to treat patients with ARVC caused by pathogenic variants in *PKP2*. The purpose of the present study was to examine the hypothesis that AAVrh.74-mediated delivery of an exogenous PKP2a gene in the setting of PKP2 deficiency in an adult mammalian heart can arrest the progression of the arrhythmogenic and cardiomyopathic components of the disease and significantly prolong life expectancy. For this purpose, we utilized a previously characterized cardiomyocyte-specific, tamoxifen (TAM)– activated, PKP2 conditional knockout murine model (PKP2-cKO).^5^ Previous studies in these animals have shown that loss of PKP2 expression in adult mice leads to an arrhythmogenic cardiomyopathy of right ventricular predominance 21 days after TAM injection, a decrease in left ventricular systolic function by 28 days post-TAM, and progression to end-stage heart failure and death by the sixth week after TAM injection. As such, this animal model presents, in a compressed time span, the various stages of human disease.^2, 5^ The purpose of the present study was to assess whether AAVrh.74-mediated delivery of PKP2a in the PKP2-cKO mouse model can arrest the progression of the arrhythmia burden and cardiomyopathy components of the disease while significantly prolonging survival. These data provide preclinical support to the notion that gene replacement therapy effectively interrupts the progression of an otherwise deadly condition, paving the way for future translation to a carefully selected human patient population that could benefit from a gene therapy approach.

## METHODS

Experiments were performed in 3-6 month old male mice expressing a cardiac-specific, tamoxifen (TAM)-activated deletion of PKP2 (PKP2-cKO)^5^. Cre-negative, flox-positive, TAM-injected age– and gender-matched mice were used as controls. Procedures conformed with the Guide for Care and Use of Laboratory Animals of the National Institutes of Health and were approved by the NYU-IACUC committee [#160726-03].

Administration of approximately 100 µL of AAV vector (corrected by body weight to reach a consistent final dose as specified in the figure legends), or Formulation Buffer (FB), was performed by a single tail vein injection. AAVrh.74-PKP2a or FB was administered through a 28– gauge sterile stainless-steel needle interfaced with a standard 0.5 mL insulin syringe (BD). Before treatment, fresh samples of AAVrh.74-PKP2a and FB were prepared to generate the appropriate concentration and maintained on wet ice.

Details related to AAVrh.74-PKP2a vector design, production and characterization as well as procedures for echocardiography, Western blot, RT-ddPCR, ddPCR immunofluorescent staining, fibrosis quantification and electrocardiography followed those previously published and are detailed in the **Data supplement.**

### Data availability

The data that support the findings of this study are presented in the article and in the **Data supplement**. Additional information can be made available from the corresponding author upon reasonable request.

### Statistical analysis

Numerical results are given as mean and standard deviation (SD). All data sets were tested for normal distribution by the Shapiro-Wilk and Kolmogorov-Smirnov tests, and significance was determined by parametric or non-parametric methods, as appropriate. The specific methods used to determine significance are specified in the figure legends. Data were analyzed using GraphPad Prism v9.2.0.

## RESULTS

Initial experiments sought to establish that AAVrh.74-mediated PKP2a expression driven by a cardiac-selective human Troponin T promoter (hTnT; schematic presented in **Supplemental Figure 1**) could prevent the ARVC phenotype in PKP2-cKO mice. All mice received a single tail vein injection of either AAVrh.74-PKP2a, at the specified dose, or formulation buffer (FB) alone. For the first set of experiments, all mice were then injected with TAM 28 days after the injection of AAVrh.74-PKP2a or FB. Five groups of male mice, 3-4 months of age were studied: 1) Control (i.e., PKP2 flox/flox Cre-negative) injected with FB; 2) PKP2-cKO (flox/flox Cre-positive) injected with FB; 3) PKP2-cKO injected with AAVrh.74-PKP2a at a dose of 3×10^13^ vg/kg; 4) PKP2-cKO injected with AAVrh.74-PKP2a at a dose of 6×10^13^ vg/kg 5) PKP2-cKO injected with 2 x 10^14^ vg/kg (**Supplemental Figure 2A**). Hearts were collected 28 days after TAM injection (i.e., 56 days after injection of either AAVrh.74-PKP2a or FB). The presence and abundance of AAV vector DNA, transgene mRNA transcripts, and PKP2 protein in the heart lysates were determined by ddPCR, RT-ddPCR and Western blot, respectively. Cumulative results are shown in **Figure 1A-C**. Examples of Western blots are presented in **Supplemental Figure 3**. The results demonstrate efficient transduction and subsequent expression of human PKP2a (hPKP2a) transgene mRNA and PKP2 protein in the heart in all AAVrh.74-PKP2a injected mice.

**Figure 1.**
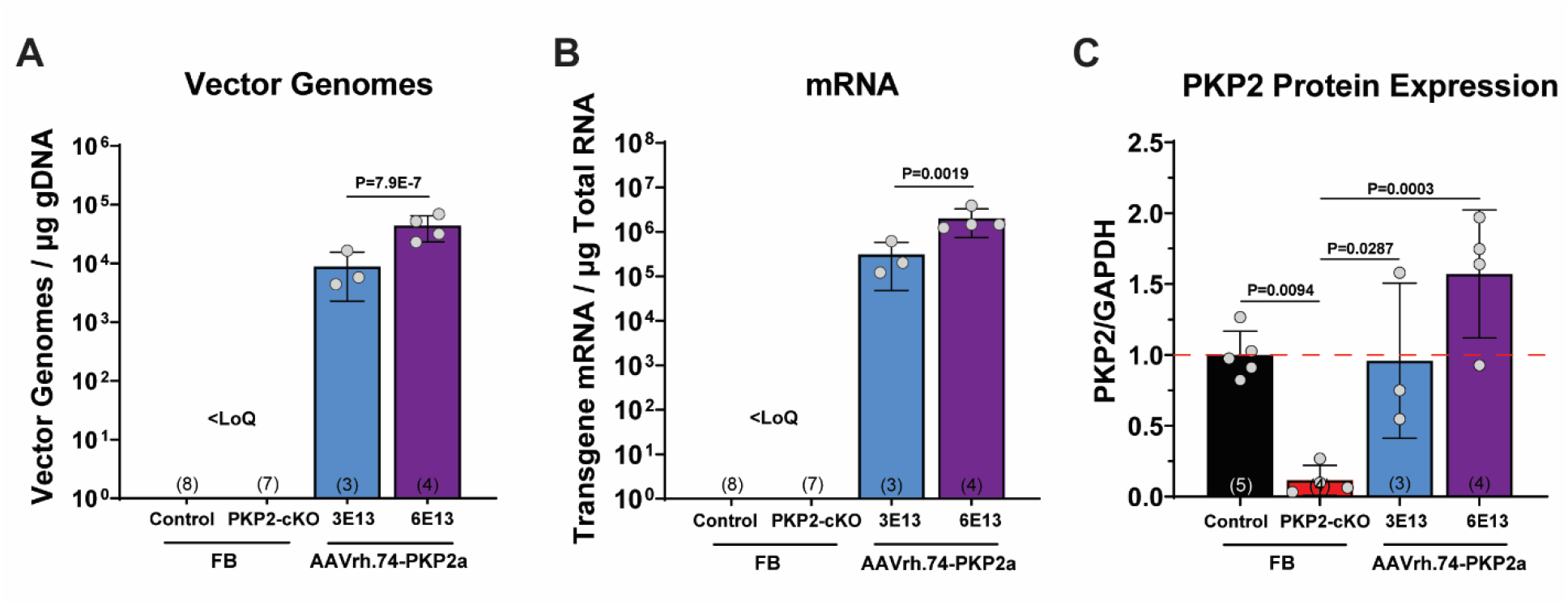
AAVrh.74-PKP2a-mediated expression in PKP2-cKO mouse heart. Dose– dependent detection of (**A**) vector genomes (ddPCR), (**B**) transgene PKP2a mRNA (RT-ddPCR) and (**C**) PKP2 protein expression (Western Blot) in cardiac tissue from AAVrh.74-PKP2a injected animals relative to controls. Red dotted line in panel C indicates endogenous expression level for PKP2 protein normalized to GAPDH in control-FB mice. Data are presented as Mean ± SD. Black bars, control mice treated with Formulation Buffer (FB); red bars, PKP2-cKO mice treated with FB; blue bars, PKP2-cKO mice treated with AAVrh.74-PKP2a 3 × 10^13^ vg/kg; purple bars, PKP2– cKO mice treated with AAVrh.74-PKP2a 6 × 10^13^ vg/kg. Number of mice studied noted in corresponding bars (N). Statistical analyses were performed using One-way ANOVA followed by Tukey’s *post-hoc* analyses. <LoQ: Less than limit of quantitation; FB: Formulation Buffer.

Immunofluorescence images acquired from fixed sections of heart tissue are shown in **Figure 2**. Panel **A** shows the immunolocalization of native PKP2 in the heart of a control (Cre-negative, TAM-injected) mouse. The image shows a clear immunoreactive PKP2-positive signal, visualized as well-defined plaques that align perpendicular to the direction of the fibers, as expected for an intercalated disc protein (see arrows in **Fig. 2A**). Panels **B** and **C** show cardiac sections of PKP2– cKO mice euthanized 28 days post-TAM and previously injected (28 days pre-TAM) either with FB (panel **B**) or with FB + AAVrh.74-PKP2a (panel **C**). Immunolabeled PKP2 signal is notably absent in the heart from PKP2-cKO mice injected with FB alone (panel **B**). However, dense PKP2 signals oriented perpendicular to the fiber orientation are present in the heart that received the AAVrh.74-PKP2a therapy (panel **C**; arrows). These results were confirmed in 4 mice per group. Overall, the data show that the exogenous PKP2 gene was transcribed and translated, and the expressed protein was properly localized to the subcellular domain expected for the native PKP2 protein.

**Figure 2.**
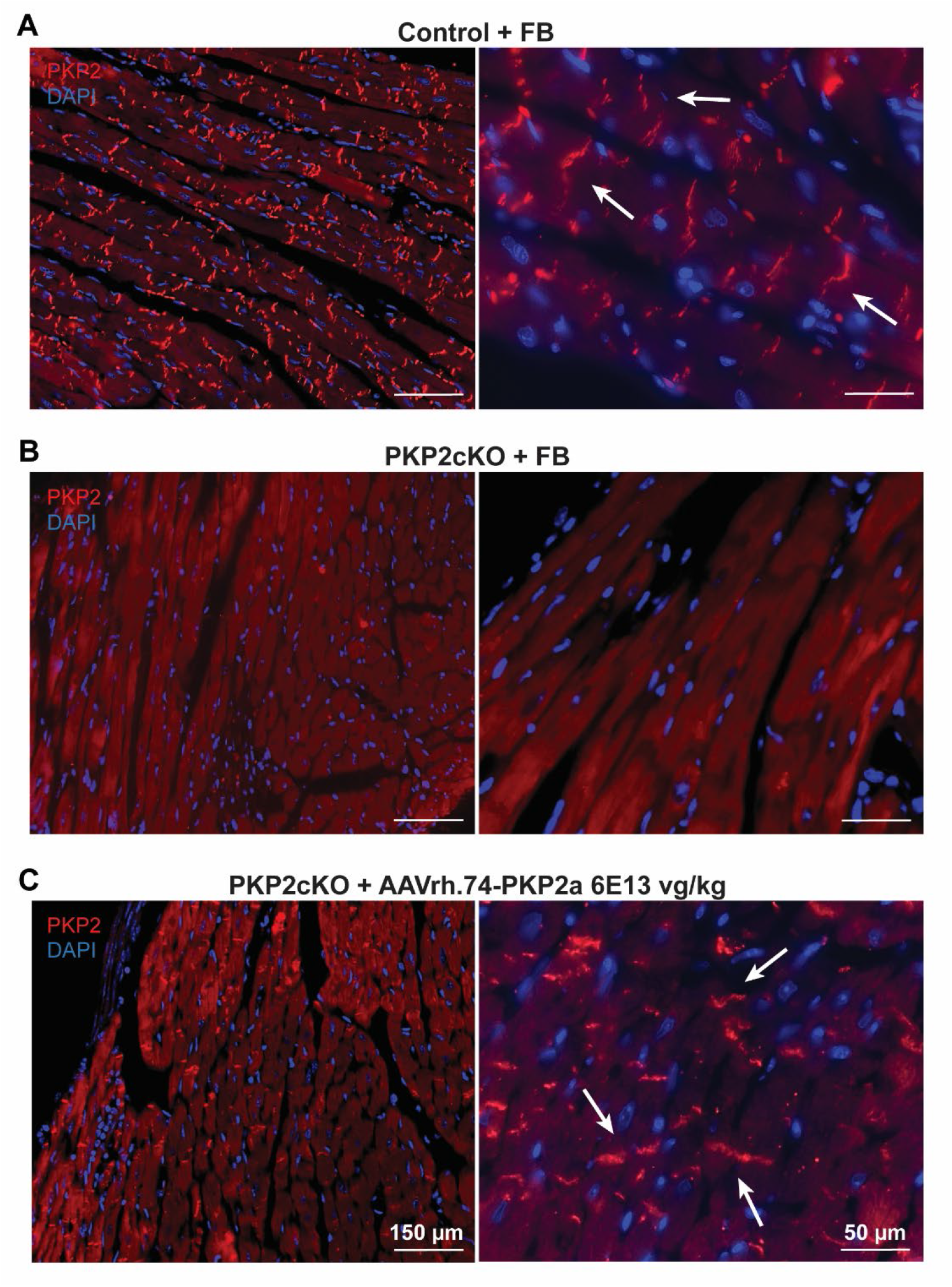
Cardiac expression of PKP2 in AAVrh.74-PKP2a treated PKP2-cKO mice. Representative images of immunofluorescence staining for plakophilin2 (PKP2; in red) and nuclei (DAPI; in blue) in hearts from (**A**) control mice treated with Formulation Buffer (FB), (**B**) PKP2– cKO mice treated with FB, and (**C**) PKP2-cKO mice treated with AAVrh.74-PKP2a 6 × 10^13^ vg/kg. Mice were treated with FB or AAVrh.74-PKP2a 28 days before Tamoxifen injection. White arrowheads in panels A and C highlight PKP2 localization at the intercalated disc. FB: Formulation Buffer. Scale bar panels on the left = 150 µm, Scale bar panels on the right = 50 µm.

We examined whether expression of the exogenous protein prevented the cardiomyopathic phenotype. Assessment of cardiac function in vector treated mice by echocardiography was performed 28-days post-TAM injection. **Figure 3** shows echocardiographic images (panel **A**) and cumulative data (panels **B-C**) obtained from hearts of mice injected with FB alone (control, black bar; PKP2-cKO, red bar), or with AAVrh.74-PKP2a (PKP2-cKO + 3×10^13^ vg/kg AAVrh.74-PKP2a, blue; PKP2-cKO + 6×10^13^ vg/kg AAVrh.74-PKP2a, purple bar). As in Figure 2, AAVrh.74-PKP2a or FB were injected 56 days before recording, and 28 days prior to TAM injection (**Supplemental Figure 2A**). As expected, loss of PKP2 expression caused a significant drop in the left ventricular ejection fraction (LVEF; red dotted line and red bar in panel B) and an increase in right ventricular (RV) area (red dotted line and red bar in panel C). In contrast, AAVrh.74-mediated expression of hPKP2a mitigated or prevented the loss of contractile function in the LV and the increase in RV area in a dose-dependent manner.

**Figure 3.**
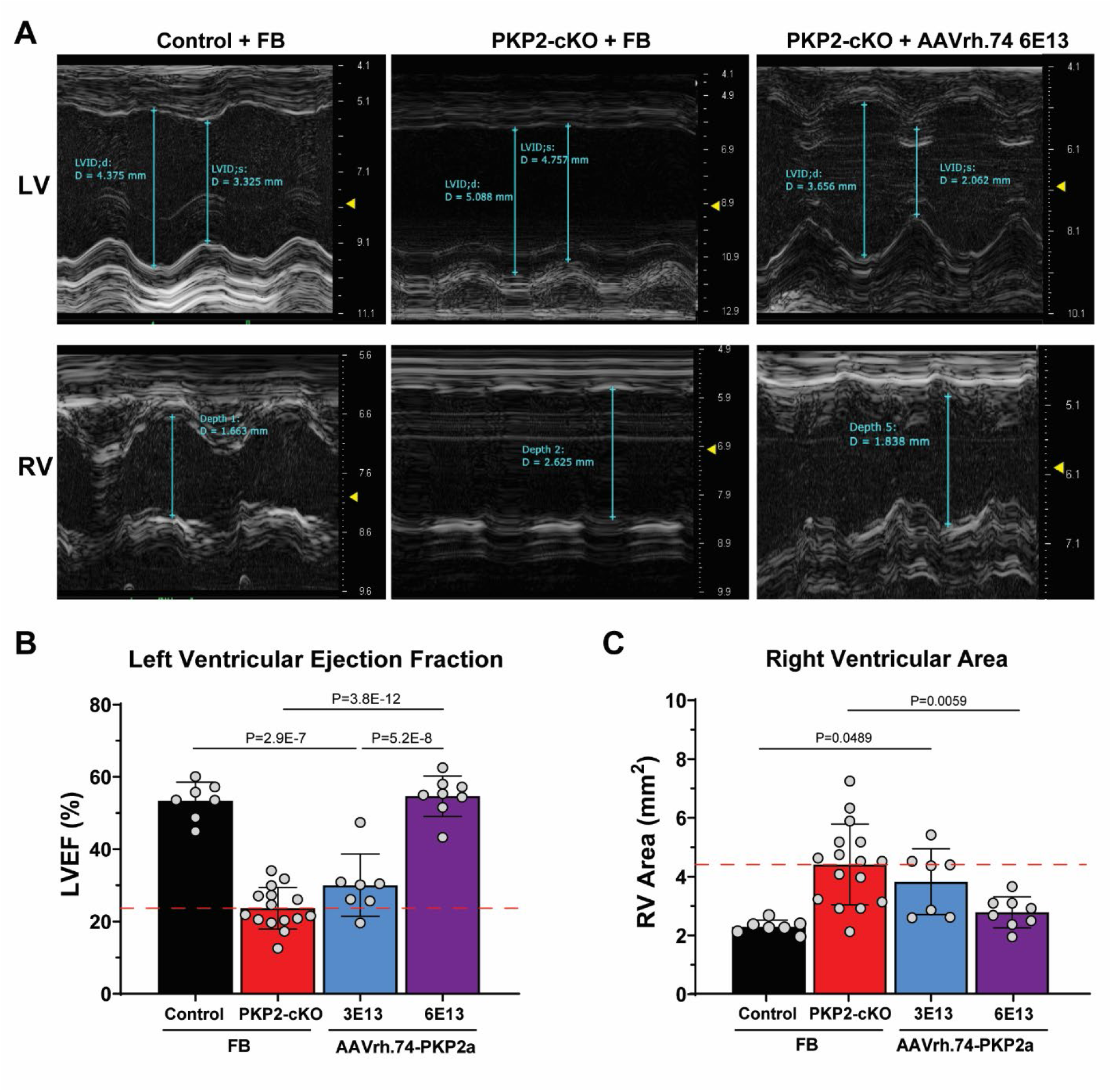
Cardiac contractility in AAVrh.74-PKP2a treated PKP2-cKO mice. (**A**) Representative images of left (LV) and right ventricular (RV) echocardiography to assess contractility in control and PKP2-cKO mice treated with Formulation Buffer (FB), or in PKP2-cKO mice treated with AAVrh.74-PKP2a 6 × 10^13^ vg/kg, 28 days before Tamoxifen injection. Measurements were collected 28 days post-Tamoxifen injection. **B)** Quantification of left ventricular ejection fraction (LVEF) measured by long axis B-mode echocardiography and **C)** right ventricular (RV) area measured by modified long axis B-mode echocardiography, as described in Cerrone et al.^5^ Red dotted lines indicate the average value recorded from PKP2-cKO mice injected with FB. Data are presented as Mean ± SD. Black bars, control mice treated with FB; red bars, PKP2-cKO mice treated with FB; blue bars, PKP2-cKO mice treated with AAVrh.74-PKP2a 3 × 10^13^ vg/kg; purple bars, PKP2-cKO mice treated with AAVrh.74-PKP2a 6 × 10^13^ vg/kg. Statistical analyses were performed using One-way ANOVA followed by Tukey’s *post-hoc* analyses. LVID: Left ventricular internal diameter, LVEF: Left ventricular ejection fraction, RV: Right ventricle, FB: Formulation Buffer.

Cardiac fibrosis is a common feature in PKP2 deficient hearts. As shown in **Figures 4A**-**C**, visualization and quantification of the extent of collagen abundance in the ventricular free walls revealed the presence of extensive fibrosis in PKP2-cKO animals (red bars), which was significantly mitigated by AAVrh.74-PKP2a administration (blue and purple bars), particularly at the higher dose (purple bar). Notably, dose-related reduction of cardiac fibrosis was observed in both the left and right ventricles of AAVrh.74-PKP2a injected animals.

**Figure 4.**
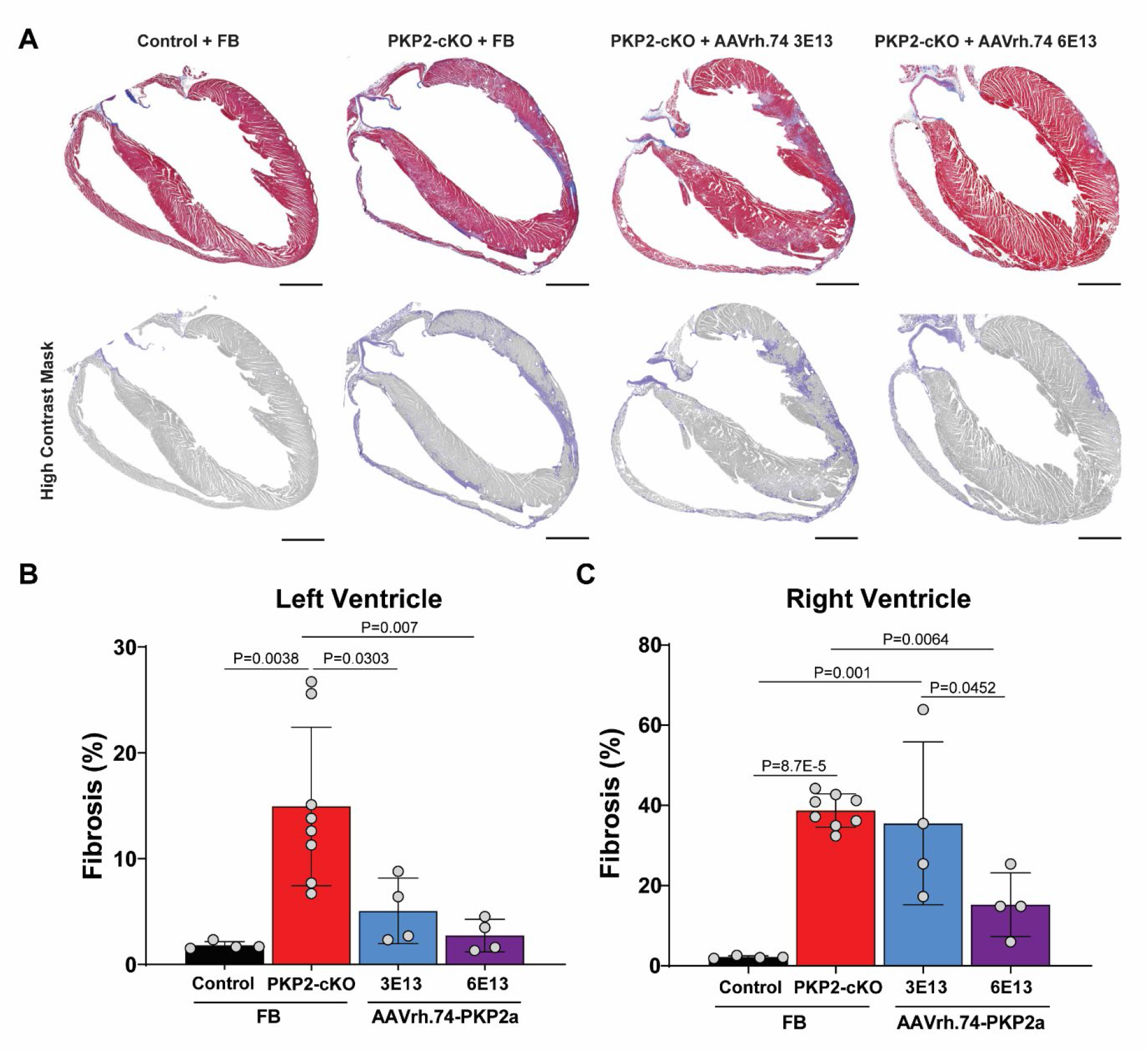
Cardiac fibrosis in AAVrh.74-PKP2a treated PKP2-cKO mice. (**A**) *Upper panel*; representative images of Masson’s trichrome staining of longitudinal heart sections of control and PKP2-cKO mice treated with Formulation Buffer (FB) and PKP2-cKO mice treated with AAVrh.74– PKP2a, 28 days before Tamoxifen injection. *Bottom panel*; High contrast mask of the same sections emphasizing collagen deposition in blue. Scale bar = 1 mm for all images. Hearts were extracted 28 days post-Tamoxifen injection. (**B & C**) Quantification of the percentage of left ventricular fibrosis (**B**) and right ventricular fibrosis (**C**) in hearts of the four different groups. Data are presented as Mean ± SD. Black bars, control mice treated with FB; red bars, PKP2-cKO mice treated with FB; blue bars, PKP2-cKO mice treated with AAVrh.74-PKP2a 3 × 10^13^ vg/kg; purple bars, PKP2-cKO mice treated with AAVrh.74-PKP2a 6 × 10^13^ vg/kg. Statistical analyses were performed using One-way ANOVA followed by Tukey’s *post-hoc* analyses. FB: Formulation Buffer.

The data presented in the previous figures demonstrated that AAVrh.74-PKP2a treatment prior to TAM-mediated knockout of PKP2 could mitigate the development of the cardiomyopathic phenotype in the PKP2-cKO mice. To evaluate whether AAVrh.74-PKP2a could arrest the progression of the ARVC phenotype when delivered after tamoxifen mediated disease induction, we studied PKP2-cKO mice injected with AAVrh.74-PKP2a 7 or 14 days **after** TAM injection (**Supplemental Figure 2B**). In addition to echocardiography analyses, the mice were followed for long-term survival. As illustrated by the Kaplan Meier curve in **Figure 5**, PKP2-cKO animals injected with FB only, died between 30 and 50 days after TAM injection, consistent with previous reports.^5^ In contrast, all but one of the mice injected with AAVrh.74-PKP2a survived for 5 months (155 days post-TAM), at which time the animals were euthanized to examine protein expression and cardiac structure. Furthermore, as shown by the representative images and cumulative data in **Figures 6A and 6B**, trichrome staining analyses revealed that the percent of the free wall of the LV and of the RV occupied by collagen in animals injected with the higher dose of AAVrh.74– PKP2a (2×10^14^ vg/kg, orange bar) was similar (though trending toward higher values) when compared to that observed in control animals injected with FB (black bar). The abundance of collagen in hearts from mice that received the lower dose of AAVrh.74-PKP2a (6×10^13^ vg/kg, purple bar) was higher than in control animals. Importantly, a comparison to collagen abundance in PKP2cKO animals at the same time point was not possible because of the early lethality in that group. However, the abundance of collagen in the AAVrh.74-PKP2a treated animals was clearly less than what was observed in PKP2-cKO mice treated with FB at 28 days post-TAM (as presented by the red dotted line in **Fig. 6B**) as well as previously published data on fibrosis in this model.^5^ Finally, cumulative data obtained from echocardiographic analysis of hearts from mice injected with AAVrh.74-PKP2a 7 days or 14 days after TAM are presented in **Figure 6C**. Hearts from FB-injected PKP2-cKO mice presented a drastic reduction in LVEF (left in **Fig. 6C**) and an increase in RV area (right in **6C**) 28 days after TAM injection (compare red bar to black bar in **Fig. 6C**), consistent with previous results.^5^ These changes were mitigated by AAVrh.74-PKP2a, even when injected 14 days after TAM, and the beneficial effects persisted up to 5 months after TAM injection, the longest time point evaluated.

**Figure 5.**
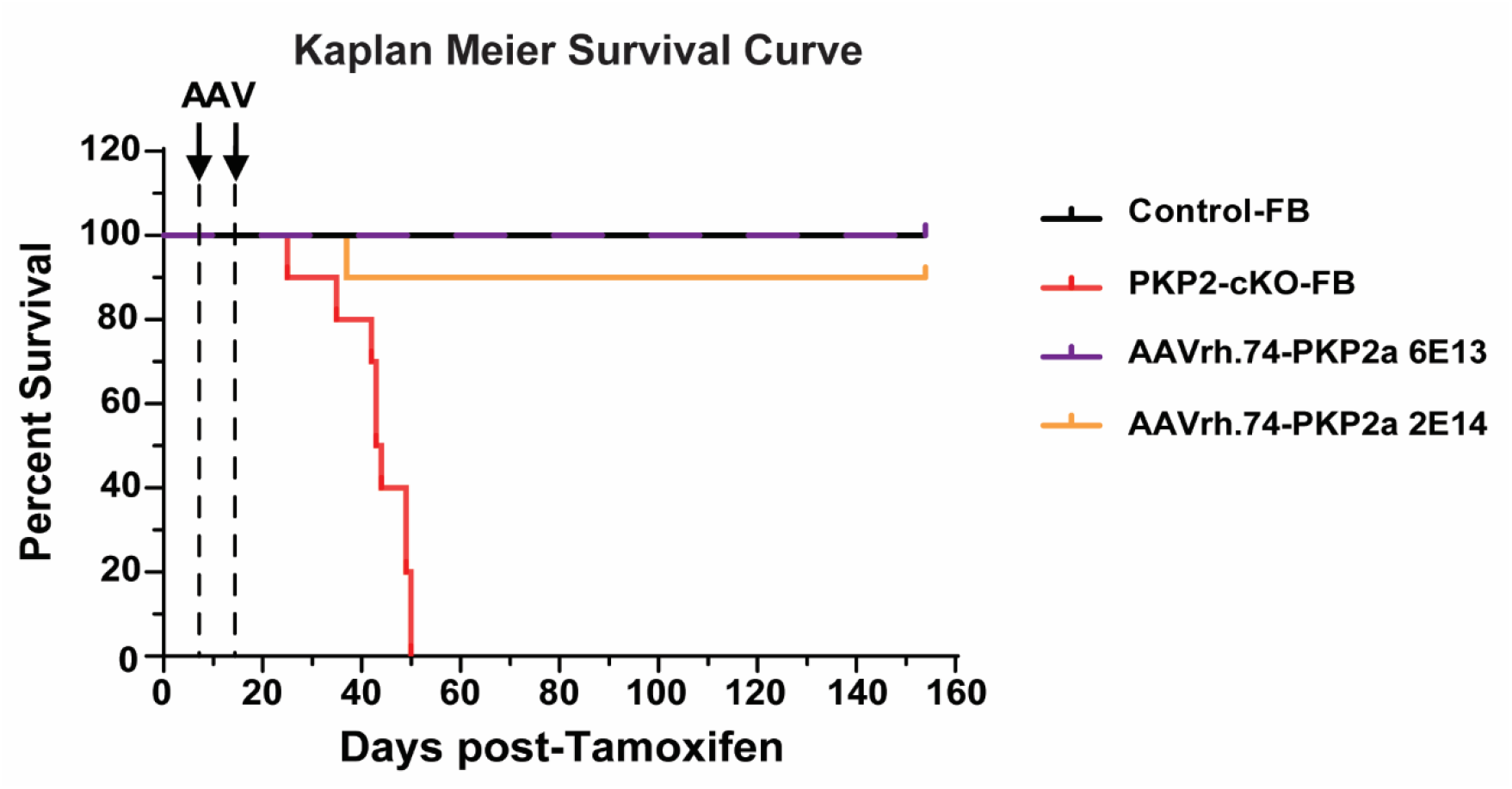
Survival following AAVrh.74-PKP2a injection in PKP2-cKO mice at 7– or 14-days post-Tamoxifen. Kaplan Meier curve depicting the long-term survival of PKP2-cKO mice following AAVrh.74-PKP2a (6 × 10^13^ vg/kg, 7 days post-Tamoxifen or 2 × 10^14^ vg/kg, 14 days post-Tamoxifen) administration compared to control and PKP2-cKO mice injected with Formulation Buffer (FB).

**Figure 6.**
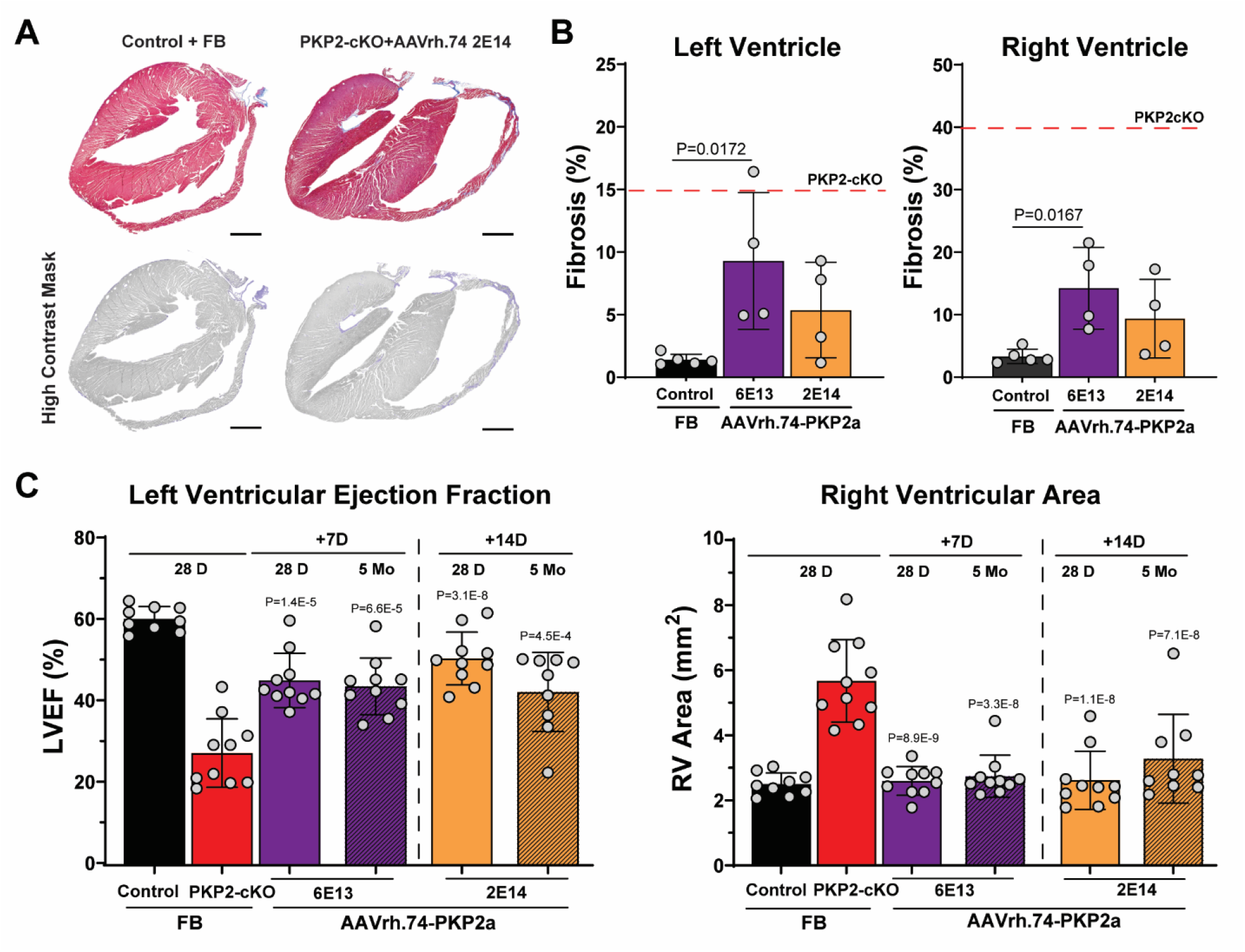
Disease progression in PKP2-cKO mice upon AAVrh.74-PKP2a treatment at 7– or 14-days post-Tamoxifen. (**A**) *Upper panel*; representative images of Masson’s trichrome staining of longitudinal heart sections of control mice treated with Formulation Buffer (FB) and PKP2-cKO mice treated with AAVrh.74-PKP2a, 2 × 10^14^ vg/kg 14 days post-Tamoxifen. Hearts were extracted 5 months post-Tamoxifen injection. *Bottom panel*; High contrast mask of the same sections emphasizing collagen deposition in blue. Scale bar = 1 mm for all images. (**B**) Quantification of the percentage of left ventricular fibrosis (left panel) and right ventricular fibrosis (right panel) by Masson’s trichrome staining of longitudinal heart sections in control mice treated with FB and PKP2-cKO mice treated with AAVrh.74-PKP2a, 7 days (6 × 10^13^ vg/kg) or 14 days (2 × 10^14^ vg/kg) post-Tamoxifen. Hearts were extracted 5 months post-Tamoxifen. Red dotted line indicates mean value of percent fibrosis recorded in PKP2-cKO mice 28 days post-Tamoxifen, injected with FB (as reported in Figure 4). (**C**) Quantification of left ventricular ejection fraction (LVEF; left panel) and right ventricular area (RV area; right panel) across time in PKP2-cKO mice treated with AAVrh.74-PKP2a, 7 or 14 days post-Tamoxifen with the dose indicated at the bottom of the bars. Echocardiography was performed at 28 days and 5 months post-Tamoxifen. Echocardiography for control and PKP2-cKO mice treated with FB was performed at 28 days post-Tamoxifen only. Data are presented as Mean ± SD. Black bars; control mice treated with FB; red bars; PKP2-cKO mice treated with FB, purple bars; PKP2-cKO mice treated with AAVrh.74– PKP2a 6 × 10^13^ vg/kg, orange bars; PKP2-cKO mice treated with AAVrh.74-PKP2a 2 × 10^14^ vg/kg. Statistical analyses were performed using One-way ANOVA followed by Tukey’s *post-hoc* analyses. FB: Formulation Buffer.

Previous studies have documented that a bolus injection of isoproterenol (ISO) 3 mg/kg leads to premature ventricular contractions in anesthetized PKP2-cKO mice 21 days after TAM injection.^5, 6^ Therefore, we used the ISO challenge protocol to determine whether AAVrh.74-PKP2a delivered 14 days after TAM can mitigate arrhythmia burden in PKP2-cKO animals. Representative ECG traces are shown in **Figure 7A**. FB-injected PKP2-cKO animals presented multiple PVCs, and 6 out of 10 animals showed more than 100 PVCs within the 30 minutes of recording after ISO injection (**Fig.7B**) with a total count of ∼300 PVCs on average (**Fig.7C**). In contrast, both parameters of arrhythmia burden were drastically reduced by administration of AAVrh.74-PKP2a at both doses tested (6 × 10^13^ vg/kg and 2 × 10^14^ vg/kg purple and orange bars, respectively, in the **Figure 7B&C**). Overall, our data show that gene therapy is a viable way of limiting the impact of loss of the native PKP2 gene on cardiac mechanical and electrical functions, with no overt toxicity at clinically applicable doses.

**Figure 7.**
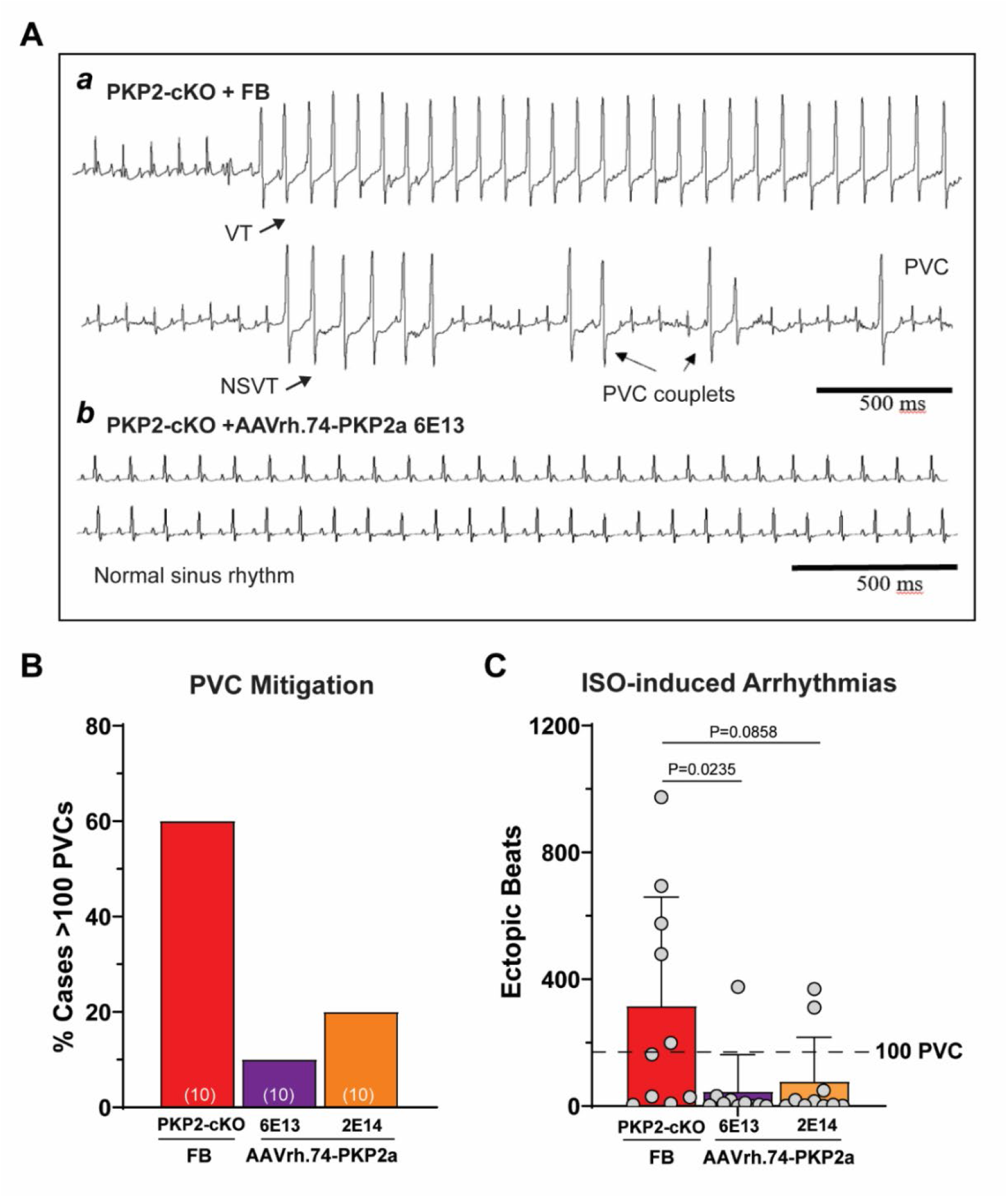
Isoproterenol-induced arrhythmias in PKP2-cKO hearts treated with AAVrh.74– PKP2a. (**A**) Representative electrocardiogram (ECG) traces from PKP2-cKO mice treated with (***a***) Formulation buffer (FB) and (***b***) AAVrh.74-PKP2a 6 × 10^13^ vg/kg, 14 days post-Tamoxifen injection. **B)** Percentage of mice that presented with >100 premature ventricular contractions (PVCs) after isoproterenol (ISO). **C**) Number of ISO-induced ectopic beats in PKP2-cKO mice treated with FB or AAVrh.74-PKP2a. Data in **B & C** was quantified over a period of 30 minutes after ISO injection and ECGs were recorded 21 days post-Tamoxifen. Data presented as Mean ± SD. Red bars, PKP2-cKO mice treated with FB; purple bars, PKP2-cKO mice treated with AAVrh.74-PKP2a 6 × 10^13^ vg/kg; orange bars, PKP2-cKO mice treated with AAVrh.74-PKP2a 2 × 10^14^ vg/kg. Statistical analyses were performed using One-way ANOVA followed by Dunn’s *post-hoc* analyses. PVC: Premature ventricular contraction, FB: Formulation Buffer.

## DISCUSSION

The present study provides experimental evidence indicating that delivery of the human PKP2 gene into cardiomyocytes following administration of an AAVrh.74-PKP2a vector by a single intravenous injection, can arrest the progression of an otherwise lethal arrhythmogenic cardiomyopathy of right ventricular predominance caused by loss of expression of the native PKP2 gene. Our results demonstrate that PKP2 gene therapy can drastically improve clinical outcomes in an animal model of ARVC. The results enable the potential for a cautious pursuit of studies that can determine the utility of PKP2-based gene therapy for human patients affected with ARVC consequent to PKP2 deficiency.

Early evidence that exogenous genes can be introduced into cardiac cells via viral particles with the purpose of affecting cardiac electrophysiology was provided by Fishman and colleagues^7^ and later, by Donahue et al.^8, 9^ Those attempts highlighted both the potential of the methodological principle and the hurdles associated with it. Bongianino et al. were first to demonstrate that AAV– mediated gene delivery can restore function in an animal model of an inheritable arrhythmia disease,^10^ a demonstration later expanded upon by Liu et al.^11^ These two proof-of-principle studies focused on very rare non-RyR2 mutations causative of Catecholaminergic polymorphic ventricular tachycardia (CPVT). Recently though, the group of Ackerman has implemented a dual method to repress expression of an endogenous –mutated– gene while expressing a gene with a native coding sequence.^12, 13^ Their experiments, using cellular models of long QT syndrome have been successful, and provide a solid and encouraging framework for future implementation of this approach in the patient population. Of note, this method is a major breakthrough for the potential of gene therapy under conditions in which the mutated protein acts as a dominant negative component that disrupts function. Clinically-relevant conditions resulting from strict loss-of– function mutations would not require repression of the native –mutated–gene for the replacement therapy approach to be successful.

Strong support for the hypothesis that AAV-mediated gene therapy may confer benefit in cardiomyopathy can be found in translational studies for Danon disease, an X-linked autophagic vacuolar myopathy, in which LAMP2B gene transfer has been found to improve metabolic and physiologic function in a preclinical animal model^14^ and has subsequently been found in clinical trials to result in confirmed transgene protein expression in the heart with improvements in key cardiac biomarker and functional measures in this patient population.^15^

In the adult mammalian heart, PKP2 is densely localized in the intercalated disc, where it integrates into structural complexes necessary for intercellular adhesion (desmosomes and area composita).^16–18^ Our data show that this precise localization is retained for the AAV-mediated expression of the PKP2 protein, thus indicating that trafficking mechanisms necessary for adequate localization of PKP2 to its functional site remain available for the exogenous protein. Although detailed future studies will be necessary, it is important to note that we did not observe diffuse or erroneous cellular localization of PKP2 in AAVrh74-injected PKP2-cKO mouse myocytes. The latter suggests that, at least at the doses injected, the cellular machinery was not overwhelmed by the presence of the exogenous gene and/or associated therapeutic hPKP2a protein expression.

Western blot analysis revealed the persistence of a light PKP2-immunoreactive band in heart lysates obtained from PKP2-cKO mice. This result, consistent with data previously published,^5^ is likely consequent to the fact that PKP2 is also expressed in non-myocyte cells in the heart such as in epicardial cells^19^. In accordance with the latter, we did not detect PKP2 in cardiac myocytes in histological sections of hearts from PKP2-cKO mice injected with formulation buffer alone, though PKP2 expression was observed in the AAV-injected animals (our Figure 2).

The murine model utilized in the present study has allowed us to learn about the importance of PKP2 in the heart, and its role in maintaining structural and functional homeostasis. Yet, we readily acknowledge that this is an imperfect model of disease. In fact, one could argue that none of the murine or cell-based models of desmosomal deficiency accurately recapitulate all aspects of ARVC. All of these models are, however, useful as entry points in the exploration of disease mechanisms and for exploration of potential therapeutic strategies. Of importance is the absence of native protein in the PKP2-cKO mice versus the clinical setting in which patients often harbor a single-allele mutation of PKP2. This raises the question of whether adding copies of the full-length gene will overcome the functional deficiency associated with the presence of a pathogenic PKP2 variant in the genome of the patient. Though a firm answer will have to await an actual clinical trial, studies in experimental models provide a reason for optimism. Indeed, there is no evidence indicating that PKP2 mutant proteins can exert dominant negative functions, and murine models in which a PKP2 mutation has been knocked-in strongly suggest that the resulting phenotype is consequent to the lack of the normal allele, and not to a dominant effect of the mutation per se.^20, 21^ Additionally, a majority of mutations underlying the clinical disorder are truncating variants^22, 23^, unlikely to generate deleterious aberrant proteins. Overall, the data suggest that PKP2-related ARVC is a loss-of-function disease and exogenous introduction of the full-length gene would have the potential to address the loss of function. Importantly, AAV-mediated delivery of the PKP2 gene in a normal murine heart did not lead to a pathogenic phenotype,^24^ thus addressing concerns about whether excessive expression of PKP2 may have adverse consequences in the treated patient population.

All limitations notwithstanding, the data presented here support the notion that gene therapy may be effective to treat PKP2-related ARVC in appropriately selected patients. It is likely that a future clinical trial will exclude patients with antibodies against the AAV serotype used to deliver the gene.^25^ In the present study we used an adeno-associated virus vector of serotype rh74 (AAVrh.74). This vector, originally isolated from rhesus monkeys, effectively transduces myocytes^26^ and, in contrast to other serotypes isolated from humans (e.g., AAV9), appears to present at a relatively low prevalence in the human population.^27^ The successful results with AAVrh.74 in the present preclinical study support its implementation in a future clinical trial in which a majority of patients are unlikely to be excluded because of prevalent anti-AAVrh.74 antibodies.

ARVC is a disease with incomplete penetrance.^2^ While significant progress has been made in risk stratification of patients suspect of ARVC,^28–30^ it remains unpredictable whether or when the disease will present in an asymptomatic gene carrier.^2^ As such, any future implementation of gene therapy in PKP2 pathogenic variant carriers will likely not be in an asymptomatic individual as a preventive measure, but as a therapeutic procedure for a patient with an overt disease. In that regard, we note that gene replacement in our model was effective even when the AAVrh.74-PKP2a was injected 14 days post-TAM, a time point at which no PKP2 can be detected in the cardiomyocytes and an increase in RV area can already be detected by echocardiography (see^5^). It is also important to highlight that reintroduction of the PKP2 gene arrested the progression of the cardiomyopathy but did not revert the functional damage. This is consistent with the notion that gene therapy in a cardiomyopathy associated with loss of muscle mass (such as ARVC) would not be expected to restore the muscle mass but only to prevent further loss as the disease progresses.

Recent studies have highlighted the importance of an inflammatory and innate immune response in PKP2-deficient hearts and myocytes.^31, 32^ We speculate that as the AAVrh.74-mediated delivery of PKP2 arrests disease progression, it may also arrest the inflammatory process, but, additional experiments will be necessary to address this important question.

The present study focused on PKP2, the desmosomal gene most commonly associated with a gene-positive arrhythmogenic cardiomyopathy of right ventricular predominance. Recent studies suggest that the disease process consequent to variants in other desmosomal genes may follow a course different from that of PKP2, making it attractive to move to a nomenclature system in which the affected gene is represented in the name of the disease (e.g., a desmoplakin cardiomyopathy^33^). Along those lines and to emphasize the heavy arrhythmogenic component that is often present in patients with PKP2 pathogenic variants, we propose to use the term plakophilin-2 arrhythmogenic cardiomyopathy (PKP2-ACM) as an entity of its own, one that, among the desmosomal cardiomyopathies, offers a feasible target for gene therapy.

In conclusion, we have conducted proof-of-principle preclinical studies to determine the potential for gene therapy in the setting of an arrhythmogenic cardiomyopathy associated with PKP2 loss of function in the murine heart. Our results indicate that a single intravenous injection of AAVrh.74-PKP2a delivering the PKP2 gene leads to the production and proper localization of PKP2 in adult cardiac myocytes, arrests the progression of the disease, and drastically shifts survival, from a condition leading to 100% lethality to 100% survival in the treated population. Our data represent one component of the necessary body of evidence to support implementation of gene therapy in.a carefully selected fraction of patients. While the treatment was well tolerated and effective in the experimental model, data from extensive preclinical safety studies are also needed to ensure safe and successful implementation in the treatment of patients in need.

## SOURCES OF FUNDING

Partly supported through a sponsored research agreement between Rocket Pharmaceuticals, Inc. and NYU Grossman School of Medicine.

## DISCLOSURES

There are no past, present or future royalty obligations for MD, MC, GC, MZ and CJMvO. BN, CBS, DR, KMS, VM, EF, PY, JS and CDH are employees of Rocket Pharmaceuticals, Inc and as such receive a salary and stock options.

## SUPPLEMENTAL MATERIAL

Supplemental Materials and Methods Supplementary Figures S1-3

## SUPPLEMENTAL MATERIAL

### DETAILED METHODS

#### Administration of AAV vector

Administration of approximately 100 µL of each AAV vector (depending on body weight), or Formulation Buffer (FB), was performed by a single tail vein injection. AAVrh.74 or FB were administered through a 28-gauge stainless steel needle interfaced with a standard 0.5 mL insulin syringe (BD). Before treatment, fresh samples of AAVrh.74-PKP2a and FB were prepared to generate the appropriate concentration and maintained on wet ice.

#### Echocardiography

Animals undergoing echocardiography (a non-invasive procedure) were lightly anesthetized with isoflurane (isoflurane in O_2_ maintained at 1-1.5% after induction). Echocardiography was performed in anesthetized animals, and body temperature was maintained at 36.5-37°C, using a Vevo 2100 machine (Visualsonics Inc) as described in Cerrone et al.^5^ Postoperative procedures were conducted in accordance with approved IACUC guidelines and involved observation until animals were fully active and recovered from anesthesia, at which time they were returned to the housing facility. Quantitative measurements were analyzed using the Vevo2100 analytical software. The left ventricular ejection fraction (LVEF, %) was calculated using B-mode parasternal long axis view and volume measures, automatically using the proprietary Visualsonics Inc. software (vsi_cardiac package.sxml). The right ventricular (RV) area was calculated using the B-Mode modified parasternal long axis view of the RV to visualize the chamber and determine the RV diameter measured halfway between the pulmonary and tricuspid valves from the interventricular septum to the free wall during diastole as in Cerrone 2017.^5^

#### Electrocardiogram recordings

Electrocardiography recordings, in combination with an isoproterenol (ISO) challenge, was performed on AAVrh.74-PKP2a and FB injected PKP2-cKO mice at 21 days post-tamoxifen injection. Mice were anesthetized with 1.5% isoflurane in 700 ml O_2_ per minute via a nose cone (following induction in a chamber containing 3-4% isoflurane in O_2_) for the duration of the procedure. Rectal temperature was monitored continuously and maintained at 37-38 °C using a heating pad. ECG (leads I, II, III) was recorded via subcutaneous needle electrodes inserted in each limb. Acquisition was initiated 1-2 min after the induction of anesthesia. After anesthesia and ECG stabilization for 1-3 min, 3 mg/kg ISO was injected intraperitoneally as a single bolus and the ECG was recorded for 30 min post injection.^5, 34^ Post-acquisition, analysis was performed on the LabChart 7.0-Pro software. The number of premature ventricular contractions (PVCs) was quantified manually, and confirmed by a second operator, in order to calculate ISO challenge arrhythmic burden.

#### Western Blot to detect PKP2 expression

Ventricular samples were homogenized in extraction buffer containing protease and phosphatase inhibitors (50 mM Tris pH 8.0, 150 mM NaCl, 0.02% Sodium azide, 1% Triton X-100, 1 mM phenylmethylsulfonyl fluoride [PMSF], 1 mM Na_3_VO_4_, 50 mM NaF and Complete Protease Inhibitor [Roche]). Protein concentration was determined using a commercially available BCA kit (Thermo Fisher) according to standard laboratory protocols. Samples were run on 4-12% precast polyacrylamide SDS gradient gels (BioRad) and transferred onto nitrocellulose membranes, (BioRad) and subsequently incubated in blocking buffer consisting of phosphate buffered saline (PBS) with Tween-20 (0.1%) and 1% nonfat dry milk. Membranes were then incubated with specific primary antibody (PKP2 rabbit polyclonal, Boster Biological Tech #PA2278, 1:2000) diluted in 1% nonfat dry milk overnight at 4°C followed by wash steps and secondary antibody (Odyssey goat anti-mouse®680RD, LI-COR # 926-68070). Antigen complexes were visualized with the Odyssey Infrared Imaging System (LI-COR). Antigen complexes were visualized, and the bands intensity quantified with the Odyssey Infrared Imaging System (LI-COR).

#### RT-ddPCR to detect transgene mRNA expression

Total RNA was isolated using Trizol Reagent (Thermofisher) followed by the MagMAX kit (Thermofisher) using the manufacturer’s protocol. Transgene quantification was done using 1-step RT-ddPCR Advanced Kit for Probes (BioRad) according to the manufacturer recommendation with 2 – 50 ng mRNA/reaction. Multiplex reactions were set up using primer sets targeting vector PKP2 and mouse Hprt. Droplets were generated using an Automated Droplet Generator followed by PX1 PCR Plate Sealer to seal the plate. The generated droplet plate was then thermal cycled on the C1000 Thermal cycler and read using the QX200 Droplet Reader. The raw data was generated using the QuantaSoft Software and the threshold set to a level 3 times the average of Mean Amplitude of Negatives in NTC (negative control) wells for the transgene probe and 2 times the average Mean Amplitude of Negatives in NTC wells for the reference (Hprt) probe. Final data is presented as Transgene Copies/µg RNA.

#### ddPCR to detect Vector DNA

Vector DNA was quantified (vg/µg gDNA) by ddPCR. Genomic DNA was recovered from the heart using the QIAamp 96 DNA QIAcube HT Kit (Cat Number 5133, Qiagen) per the manufacturer’s instructions. ddPCR multiplexed reactions were set up using the ddPCR Supermix for Probes (No dUTP) from Bio-Rad according to the kit’s instructions with genomic DNA and primers/probe set targeting vector PKP2. A mouse Tfrc probe was used as the housekeeping gene to normalize the vector DNA copy numbers per host gDNA. Droplets were generated using the Auto Droplet Generator (Bio-Rad) and subjected to PCR reaction using the C1000 Thermal Cycler. The droplets were read using the QX200 Droplet Reader all according to manufacturer’s instructions. The raw data was generated using the QuantaSoft Software and the threshold set to a level 3 times the average of Mean Amplitude of Negatives in NTC (negative control) wells.

#### Immunofluorescent labelling of PKP2

Hearts were fixed in 4% paraformaldehyde (PFA) overnight, then embedded in paraffin, sectioned (5 μm) and collected on Superfrost Plus microscope slides (Fisher Scientific). Sections were deparaffinized at 60 °C for 30 min, cooled to room temperature, and rehydrated by multiple sequential immersions in Xylene, 100% Ethanol, 95% Ethanol and 70% Ethanol followed by multiple washes with 1X PBS. Antigen Retrieval was performed for 30 min in 1X citrate buffer, and the slides were cooled and washed in 1X PBS. Blocking was performed for 1 hr at room temperature in blocking buffer [1X PBS containing 2% bovine serum albumin, 2% glycine and 0.2% gelatin]. Sections were incubated overnight at 4°C with the primary antibody (PKP2 rabbit polyclonal, Boster Biological Tech #PA2278, 1:200) diluted in blocking buffer, washed 3 times in 1X PBS and incubated with the secondary (anti-rabbit Alexa568, Invitrogen # A-11011, 1:300) in blocking buffer in the dark at room temperature. After a 5 min wash in PBS, the sections were counterstained with DAPI and mounted with Prolong Gold. Images were taken with a BZ-X800E All-in-One Fluorescence Microscope (Keyence).

#### Masson’s Trichrome staining

Hearts were fixed with 4% PFA in PBS, embedded in paraffin, and cut into 5 μm thick sections. Sections were stained with Masson’s Trichrome according to the manufacturer’s instructions (Plysciences 25088). Briefly, paraffin embedded sections were heated at 60 °C for 30 min and rehydrated. Slides were then incubated in Bouin’s solution, 60 °C for 1 hr followed by wash steps and immersed in Weigert’s working hematoxylin solution for 10 min. After repeated washes, slides were immersed in Biebrich Scarlet–Acid Fuchsin solution for 5 min and phosphomolybdic acid solution for 10 min prior to Aniline blue incubation for 5 min. After a final wash, slides were incubated in 1% acetic acid for 1 min, rinsed, dehydrated and mounted with Permount Mounting Medium. Stained sections were scanned at a 10× magnification using a Keyence All-in-One Fluorescence Microscope (BZ-X800E, Keyence, NJ, USA). The ImageJ (NIH) software was used for analysis of tissue sections, as in Cerrone et al.^5^ Six to seven regions of interest (ROIs) for each ventricle were identified and the interventricular septum was excluded. Percentage Fibrosis was measured as the area of collagen (blue staining) normalized to the area of tissue in the ROI.

#### Data availability

The data that support the findings of this study are presented in the article and online supplementary files. Additional information can be made available from the corresponding author upon reasonable request.

#### Statistical analysis

Numerical results are given as mean and standard deviation (SD). All data sets were tested for normal distribution by the Shapiro-Wilk and Kolmogorov-Smirnov tests, and significance was determined by parametric or non-parametric methods, as appropriate. The specific methods used to determine significance are specified in the figure legends. Data were analyzed using GraphPad Prizm v9.2.0.

## SUPPLEMENTAL FIGURES

**Supplemental Figure 1.**
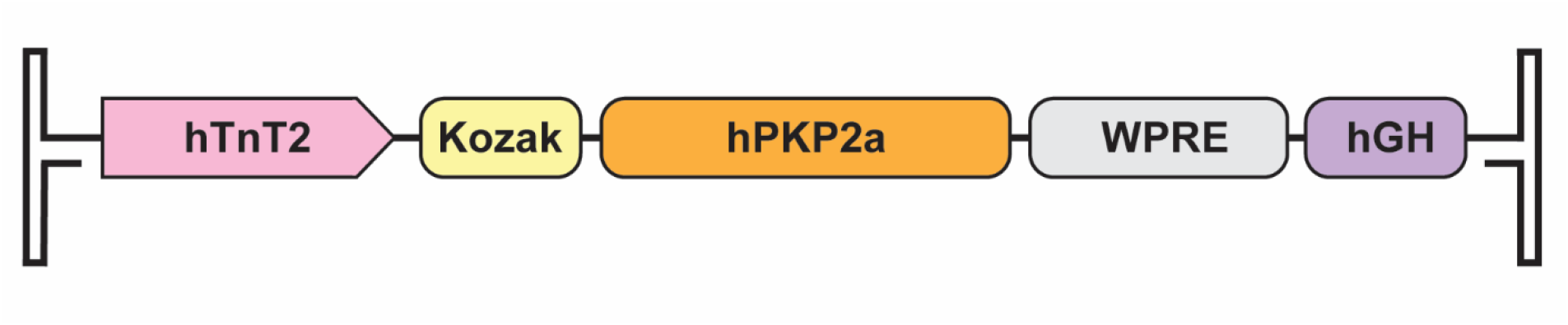
Schematic representation of the AAVrh.74-PKP2a expression cassette. The vector contains an expression cassette flanked by inverted terminal repeat (ITR) elements derived from AAV2, with the human Troponin T (hTNNT2; hTnT) promoter, consensus Kozak sequence, PKP2a transgene, woodchuck hepatitis virus post-transcriptional regulatory element (WPRE), and the human growth hormone polyadenylation signal (hGH).

**Supplemental Figure 2.**
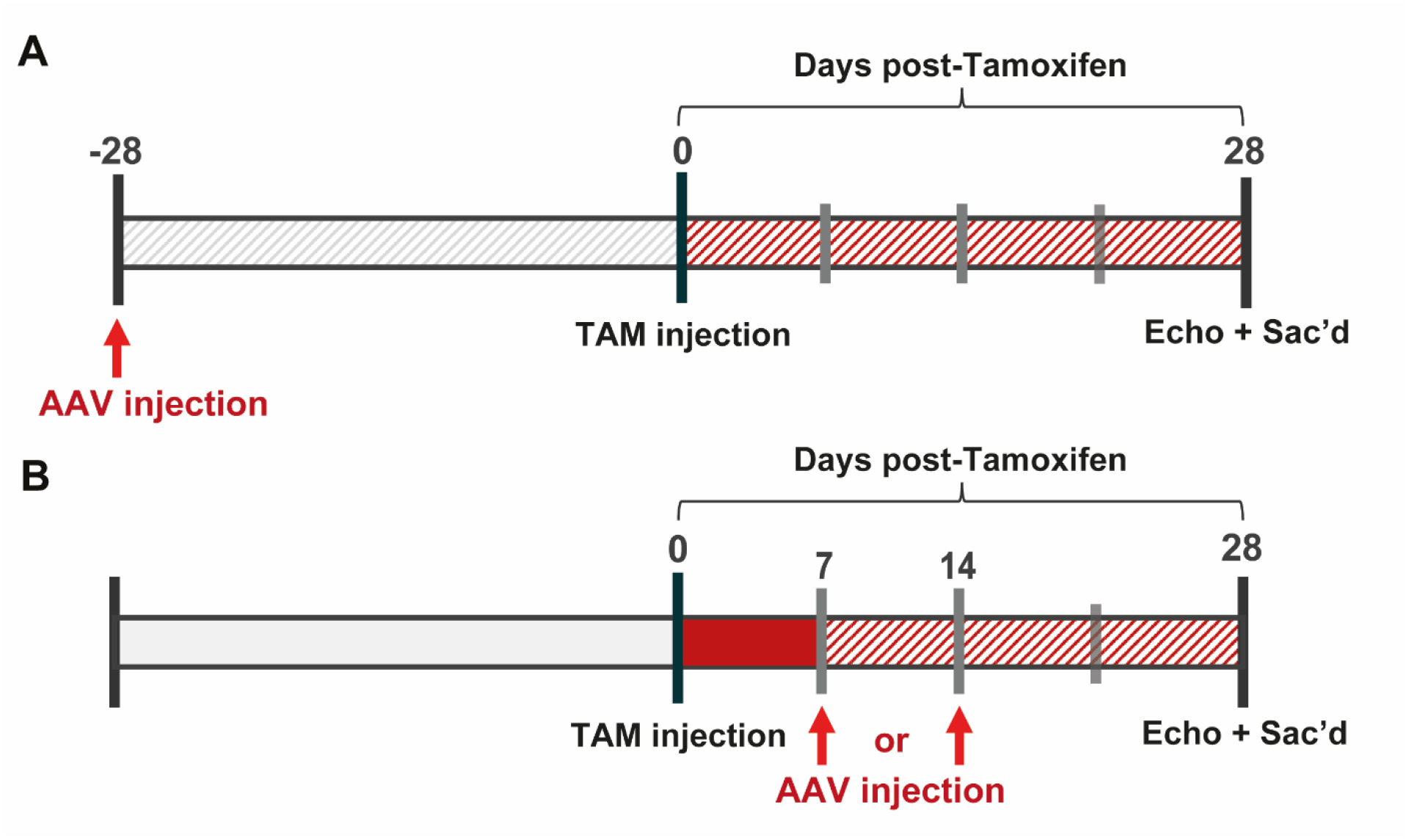
Timeline of AAVrh.74-PKP2a treatment in PKP2cKO mice. AAV treatment was delivered at **A**) 28 days before Tamoxifen (TAM) injection or **B**) 7 or 14 days post-TAM injection.

**Supplemental Figure 3.**
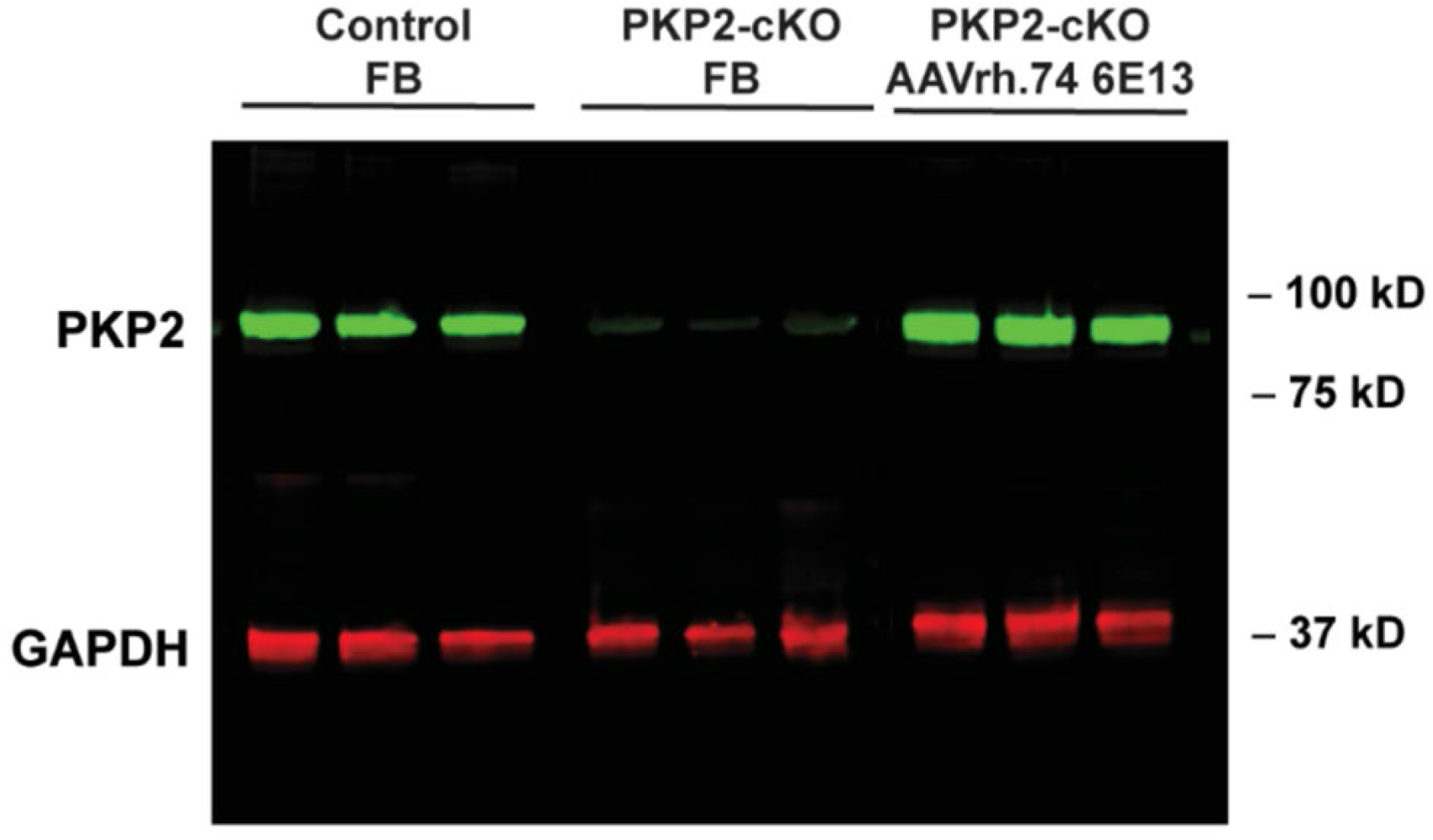
Cardiac Plakophilin 2 protein levels in PKP2-cKO by Western Blot. Gel images of PKP2 protein detected in control hearts from mice injected with Formulation Buffer (FB), hearts from PKP2-cKO mice injected with FB and hearts from PKP2-cKO mice injected with AAVrh.74-PKP2a 6 × 10^13^ vg/kg. Molecular weight of PKP2 is 97.4 kDa.

